# Lactate receptor HCAR1 regulates neurogenesis and microglia activation after neonatal hypoxia-ischemia

**DOI:** 10.1101/2020.12.02.408070

**Authors:** Lauritz H. Kennedy, Emilie R. Glesaaen, Vuk Palibrk, Marco Pannone, Wei Wang, Ali H.J. Al-Jabri, Rajikala Suganthan, Niklas Meyer, Xiaolin Lin, Linda H. Bergersen, Magnar Bjørås, Johanne E. Rinholm

## Abstract

Neonatal cerebral hypoxia-ischemia (HI) is the leading cause of death and disability in newborns with the only current treatment option being hypothermia. An increased understanding of the pathways that facilitate tissue repair after HI can aid the development of better treatments. Here we have studied the role of lactate receptor HCAR1 (Hydroxycarboxylic acid receptor 1) in tissue repair after HI in mice. We show that HCAR1 knockout (KO) mice have reduced tissue regeneration compared with wildtype (WT) mice. Further, proliferation of neural progenitor cells and microglial activation were impaired after HI. Transcriptome analysis showed a strong transcriptional response to HI in the subventricular zone of WT mice involving about 7300 genes. In contrast, the HCAR1 KO mice showed a very modest response to HI, involving about 750 genes. Notably, fundamental processes involved in tissue repair such as cell cycle and innate immunity were dysregulated in HCAR1 KO. Taken together, our data suggest that HCAR1 is a key transcriptional regulator of the pathways that promote tissue regeneration after HI. Thus, HCAR1 could be a promising therapeutic target in the treatment of neonatal HI and other forms of brain ischemia.

## Introduction

Cerebral hypoxia-ischemia (HI) affects around 1.5 per 1000 live born births in the developed countries^1^. It is characterised by an insufficient supply of blood and oxygen to the brain, leading to cell death and brain tissue damage. Hypothermia is the mainstay of today’s treatment^2^, but a high percentage of survivors still experience long-term neurological effects, including cerebral palsy, epilepsy and cognitive disabilities^3^. Following a hypoxic-ischemic event, the neonatal brain has the ability to partly regenerate^4^. This process of brain tissue regeneration requires a coordinated increase in microglia-induced inflammation, stem cell proliferation and angiogenesis. After the acute phase of cerebral HI, debris from dead cells activates microglia, the resident immune cells of the brain. These cells have the ability to remove cell debris by phagocytosis and release factors that stimulate tissue repair^5^. At the same time, the hypoxic-ischemic event leads to the release of various growth factors, which stimulate proliferation of neural stem- and progenitor cells^4^ as well as angiogenesis^4,6^. In mammals, the majority of stem- and progenitor cells are located in two proliferating areas of the brain, namely the subventricular zone, which is located adjacent to the lateral ventricles, and the dentate gyrus of the hippocampus. Proliferating cells migrate from these areas to repopulate the damaged tissue. Targeting these areas is therefore a potential strategy to stimulate repair processes after an ischemic insult.

Recent studies in mice have shown improved recovery after neonatal HI by administration of lactate^7,8^, but the underlying mechanisms for this beneficial effect are unclear. It is unknown whether lactate improves recovery by giving metabolic support to the cells or by signalling via the Hydroxycarboxylic acid receptor 1 (HCAR1), or both.

HCAR1 is a G_i_-protein coupled receptor. It was first described in adipose tissue where it inhibits lipolysis through lowering of cyclic adenine monophosphate (cAMP)^9^. In the brain, HCAR1 activation can modulate neuronal firing rates in vitro^10^ and stimulate brain angiogenesis in vivo^11^. Until now, a role of HCAR1 in tissue protection or repair after ischemia has not been demonstrated, although studies from different cell lines suggest that it may regulate cell proliferation and differentiation^12,13^.

Here we investigate the role of HCAR1 in neonatal HI in mice. We show that HCAR1 knockout (KO) mice have a reduced ability to regenerate brain tissue after HI. By examination of neurosphere cultures in vitro and immunohistochemical staining of brain tissue after HI, we find that HCAR1 KO mice display impaired proliferation of neural stem-progenitor cells and microglia. In addition, we find that microglia are less activated. Transcriptome analysis revealed that subventricular zones from HCAR1 KO mice have an almost complete lack of transcriptional response to HI. This was specific to the subventricular zone as hippocampal samples from HCAR1 KO mice responded similar to that of WT mice. Thus, HCAR1 is a crucial transcriptional regulator of tissue response to ischemia in the subventricular zone. HCAR1 could therefore be targeted to promote tissue repair after HI.

## Materials and methods

### Animals

HCAR1 KO and C57Bl/6N (WT) mice were used for this study. The HCAR1 KO line was a gift from Prof. Dr Stefan Offermanns, Max Planck Institute for Heart and Lung Research, Bad Nauheim, Germany and has been described previously^9^. The KO line was maintained in C57Bl/6N background, and genotypes were determined by PCR analysis with DNA samples extracted from mouse ears. All mice were housed in a climate-controlled environment on a 12h light/dark cycle with free access to rodent food and water. All efforts were made to reduce the number of animals used in experiments. Both females and males were included in the analyses. The mice were treated in accordance with the national and regional ethical guidelines and the European Union’s Directive 86/609/EEC. Experiments were performed by FELASA-certified personnel and approved by the Norwegian Animal Research Authority.

### Mouse model for cerebral HI

Cerebral hypoxia and ischemia (HI) was produced in P9 mice by permanent occlusion of the left common carotid artery (CCA) followed by systemic hypoxia, as previously described^14^. In brief, pups were anaesthetised with isoflurane (4% induction in chamber, 2.5% maintenance on mask in a 2:1 mixture of ambient air and oxygen), and a skin incision was made in the anterior midline of the neck. The left CCA was exposed by blunt dissection and carefully separated from adjacent tissue. A needle was placed the CAA, and a monopolar cauteriser (Hyfrecator 2000; ConMed) at a power setting of 4.0 W was used to electrocoagulate the artery. The neck incision was closed with absorbable sutures (Safil 8–0 DRM6; B. Braun Melsungen AG). The surgical procedure was completed within 5 min. After a recovery period of 1–2 h, the pups were exposed to a hypoxic (10% oxygen balance nitrogen; Yara), humidified atmosphere for 60 min at 36.0 °C. The pups were returned to their dam and after 6 hours, 24 hours or 42 days brains were retrieved and prepared for immunohistochemistry, cell culture experiments, or RNA sequencing.

### Measurement of acute tissue damage and long-term tissue loss

Mice were terminated by neck dislocation 24 h or 42 d after HI. Brains were removed from the skull and freed from dura mater and vascular tissue before being transferred to a precooled brain mould immersed in ice-cold PBS. 1□mm coronal slices were cut using an adult brain slicer (51-4984; Zivic Instruments, Pittsburgh, PA, USA). For measurement of acute tissue damage, sections were soaked in 2% TTC (T8877, Sigma) in PBS for 30□min at room temperature and subsequently fixed in 4% paraformaldehyde (PFA, Sigma-Aldrich, St. Louis, MO, USA; 15,812-7) in PBS at 4□°C for 1 hour. Photos were captured with a digital camera (Nikon D80), and pictures were analysed using image J software (NIH, San Francisco, CA, USA). Quantification of acute tissue damage/infarct size was carried out as previously described^31^. Briefly, the infarct area was calculated by subtracting the area of undamaged, TTC positive tissue in the ipsilateral hemisphere from that of the intact contralateral hemisphere, thereby correcting for brain oedema. The relative size of the damage was expressed as per cent of the contralateral hemisphere. Total volume loss, as well as tissue loss within specific brain structures, was calculated by modified Cavalieri’s principle, using the formula V=∑APt where V is the total volume, ∑A is the sum of the areas measured, P is the inverse of the section sampling fraction and t is the section thickness. For measurement of long-term tissue loss 42 days after HI, coronal sections were prepared as described above, but without TTC staining. Sections were fixed in 4% paraformaldehyde (PFA) in PBS for 30□min, and 10% formalin for 24□h and photos were taken. Tissue loss was calculated by subtracting the total volume, section volume or structure volume of the ipsilateral hemisphere from that of the contralateral hemisphere. One brain (HCAR1 KO) was excluded from the analysis due to so excessive damage that the sections fell apart, thus making measurements difficult. The person performing measurements was blinded to genotype during the measurements.

### Neurosphere cultures and assays

Neurospheres derived from forebrains of C57BL/6 and C57BL/6 HCAR1 knockout mice at postnatal day 3 were generated and propagated as previously described with modifications^32^. Briefly, dissected brain tissues were finely minced and cultured with proliferation medium of DMEM/F12 (Invitrogen) supplemented with 2 mM glutaMax, 20 ng/ml EGF (R&D Systems), 10 ng/ml bFGF (R&D Systems), N2 supplement (ThermoFisher Scientific), B27 supplement without vitamin A (ThermoFisher Scientific), and penicillin/streptomycin. Under the proliferating condition, cells were grown as free-floating neurospheres. After 7 days in culture, cells in primary neurospheres were trypsinised with trypsin-EDTA (Invitrogen), dissociated mechanically, and placed onto 75 cm2 flasks and the neurospheres were passaged every 7 days. For neurosphere self-renewal, dissociated single NSPCs were plated at a density of 1.0□×□10^4^ per well onto 6-well suspension plates with proliferation medium. After 10 days in culture, images of the entire well were captured with EVOS microscope. The pictures were analysed using the ImageJ software to obtain the total number and average size of neurospheres per well. For the neurosphere differentiation, dissociated single NSPCs were plated at a density of 5 × 10^4^ cells per centimetre square onto tissue culture plates pre-coated with poly-D-lysine (Sigma). Cells were cultured in differentiation medium (proliferation medium minus EGF and bFGF) for 5 days with half medium changes daily, and the cells were fixed at different time points for further experiments.

### Immunocytochemistry and quantification of neurospheres

Immunocytochemistry was performed as described previously^32^. Differentiated NSPCs were fixed with 4% paraformaldehyde and treated with 0.1% Triton X-100/PBS. Following blocking with 5% BSA, 5% goat serum, and 0.1 Triton X-100 in PBS for 30 min, the cells were incubated with monoclonal anti-neuron-specific beta-III tubulin (Tuj-1, MAB1195 R&D Systems), rabbit polyclonal astrocyte-specific anti-glial fibrillary acidic protein (GFAP, Z0334 Dako) in PBS containing 0.5% BSA, 0.5% goat serum, and 0.1% Tween 20 at 4°C overnight. Then the cells were incubated with fluorescent anti-mouse or rabbit secondary antibody (Alexa 594 and Alexa 488, Molecular Probes). The nuclear dye 4’,6-diamidino-2-phenylindole (DAPI) at 1 l ng/ml (Molecular Probes) was added to visualise all cells. To obtain the percentage of each cell type, 4000 –5000 cells that were morphologically identified in 10 random fields from two different cultures were counted under a 10x objective. Percentage of positive cells was calculated in relation to the total number of cells, as detected by DAPI nuclear staining.

### BrdU incorporation in mice

For the BrdU experiments, we injected 0.1□mg/g of BrdU into the peritoneum at day 4 to 7 after hypoxic ischemia at 24-h intervals. Animals were transcardially perfused 2□h after the last injection and brains were fixed and stained as described below.

### Preparation of mouse tissue and immunolabelling

Mice were anesthetised with a cocktail consisting of Zoletil Forte (Virbac International, Carros Cedex, France), Rompun (Bayer Animal Health GmbH, Leverkusen, Germany) and Leptanal (Janssen-Cilag International NV, Beerse, Belgium). Mice were then transcardially perfused with 4% PFA. Brains were then removed and stored in 4% PFA for 24□h, and then immersion fixed in 10% formalin until paraffin embedding. We then cut 6-8-μm-thick coronal sections through the entire forebrain using a microtome (ThermoScientific, Waltham, MA, USA). For immunostaining, sections were heated in an incubator at 60°C for 30 min. Deparaffinisation/dehydration was performed by immersing in Neoclear (2x 5 min, Millipore, Darmstadt, Germany) followed by rehydration in an EtOH gradient (100% 2 × 5□min; 96% 5□min and 70% 5□min) and then transferred to MQ H_2_O. Sections were then incubated at 100□°C in citrate antigen retrieval buffer (pH 6.0) for 20 min using a coverslip-paperclip method described by Vinod et al^33^. Following antigen retrieval, slides were incubated in blocking solution (10% normal goat serum, 1% bovine serum albumin, 0.5% Triton X-100 in PBS) for 1 h. Primary antibody incubation was done at room temperature overnight. Primary and secondary antibodies were diluted in a solution containing 3% normal goat serum, 1% bovine serum albumin, 0.5% Triton X-100 in PBS. The next day, sections were washed 3 x 10 min in PBS and then incubated with secondary antibodies for 1h at room temperature before a new wash of 2 × 10 min in PBS and a third incubation with DAPI for 15 minutes. Sections were rewashed (3 × 10 min) before being mounted with ProLongTM Glass Antifade Mountant (Fisher Scientific, Waltham, Massachusetts). Cover glass thickness was 0.13-0.17 mm. Primary antibodies were rat anti-BrdU (AB6326 Abcam, 1:200), guinea pig anti-Doublecortin (AB2253 Abcam, 1:500), rabbit anti-Ki67 (ab15580 Abcam) and rabbit anti-Iba1 (019-19741 Wako, 1:500). Secondary antibodies used were Alexa 488 goat anti-rabbit, Alexa 555 goat anti-rat, Alexa goat anti-guinea pig (all diluted 1:400).

### Analysis of immunolabelling

Images were captured on a Leica SP8 confocal microscope, using a 20x objective (n.a. 0.75, microglia experiments) and a 40x oil-immersion objective (n.a. 1.3, DCX experiments). All image analysis was done in Fiji ImageJ. Before analysing, Z-stacks were flattened with maximum z-projection. For the doublecortin analysis (Fig. 2), we analysed four images per hemisphere from two different sections. The subventricular zone was defined as the area within 0-50 μm from the ependyma and the intermediate zone as 50-200 μm. We counted cells manually using the Cell Counter plugin in Fiji Image J with operators blinded during imaging and analysis. In the microglial experiments, the blinded operators defined the ischemic core and peri-infarct zone by the morphological appearance of microglia, cell-cores and general cyto-architectural integrity. In the microglia analysis (Fig. 3), we used the WEKA-segmentation computer learning algorithm^34^ for image segmentation. After computer training, the images were automatically segmented using the trained algorithms (model files available at request to authors). The cells were then analysed automatically utilising the Analyze Particle tool with scripts written in ImageJ (Scripts available at request to authors). In the microglia experiment, we analysed two images per hemisphere from two different sections. For microglia counting, we counted Iba1+ overlapping with DAPI+ objects, and then triple overlap with BrdU for counting of newly made microglia. Based on previous literature on microglia morphology during cerebral ischemia^19^, we selected the average maximum branch length and microglial cross-sectional size as data for activated microglia morphology analyses. For the branch and size analyses, we excluded processes protruding from out of focus microglia by only analysing Iba1+ objects above 95um^2^. Branches were analysed using the skeletonise (2D/3D) and analyse skeleton (2D/3D) functions in ImageJ.

### RNA sequencing and analysis

Subventricular zone tissue was dissected from the ipsilateral (damaged) and contralateral (undamaged) hemisphere of mice 3 days after HI. The samples were snap-frozen in liquid N_2_ and stored at −80°C before RNA isolation. RNA isolation was performed using QIAGEN allprep kit, and final RNA was dissolved in MQ H_2_O and stored at −80°C. Paired-end sequencing was performed with the Illumina platform by BGI Tech Solutions (Hong Kong). The quality control of fastq files was performed with FastQC v0.11.9^35^. Alignment to reference genome (GRCm38) was accomplished with hisat2 v2.1.0^36^ while annotation and count matrix was completed with featureCounts v.2.0.0^37^. We performed downstream DEGs analysis in R v3.6.1 with DESeq2 v1.24.0^38^. GSEA (Gene Set Enrichment Analysis) was done with WebGestalt^39^. Heat maps were generated in R v3.6.1 with heatmap3 v1.1.7^40^.

### Statistical Analysis

P-values in Fig. 1 and Fig 2G-I are from unpaired, two-tailed, t-test’s. In Fig. 2O-R and Fig. 3 we used the Šídak (Šídak-Bonferroni)^41^ method for multiple comparisons of selected groups (WT-contra vs KO-contra, WT-ipsi vs KO-ipsi, WT-contra vs WT-ipsi, and KO-contra vs KO-ipsi. Alalyses were performed in Prism. Degrees of freedom are written as df in the results. All error bars represent the standard deviation. In Fig. 2 O-R and Fig. 3, there are no error-bars as all the individual data points are shown in the graphs. All experimental units were included in the analyses (none were excluded), unless otherwise stated.

**Figure 1:**
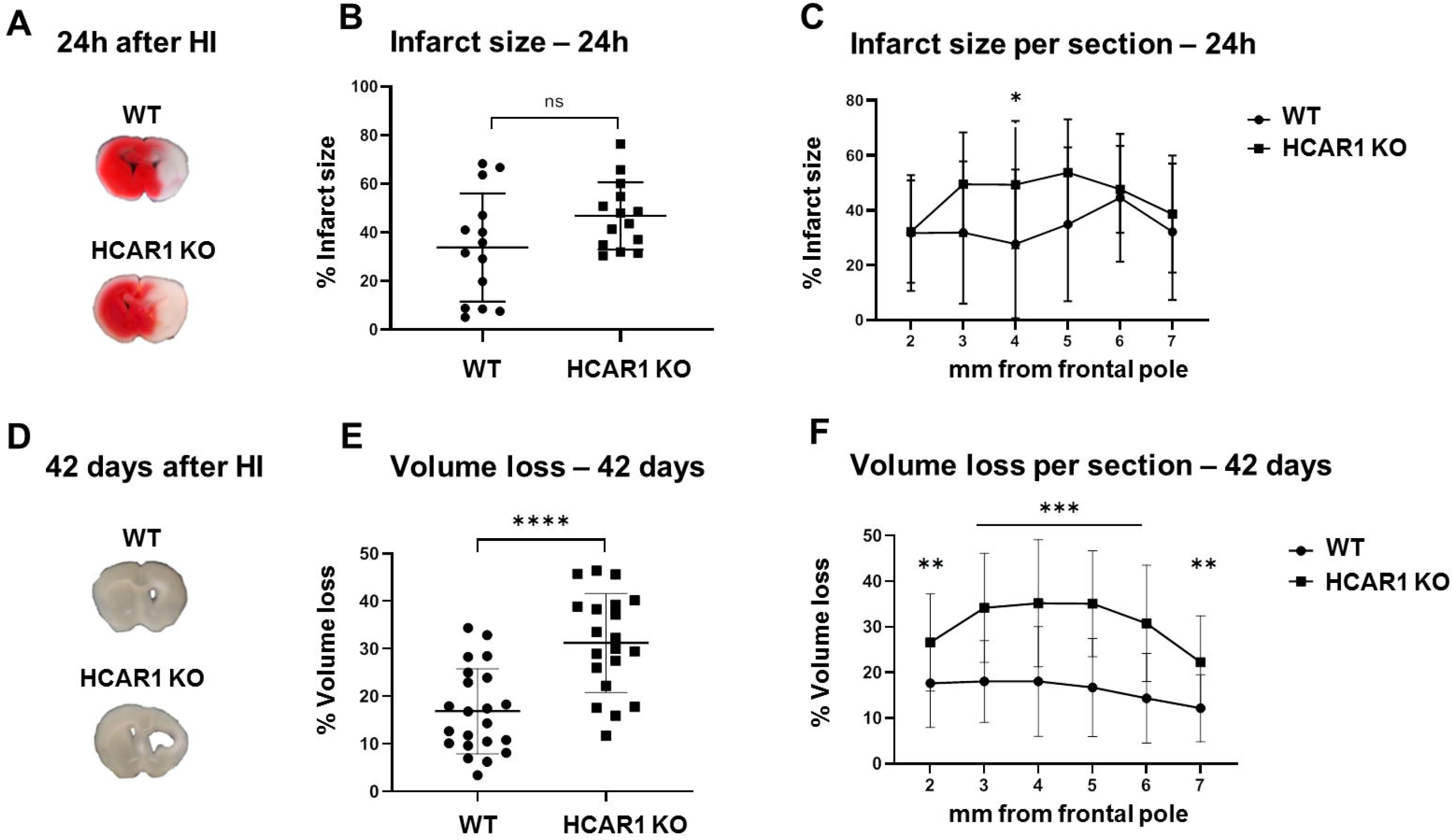
HCAR1 does not protect the brain from acute tissue damage following cerebral HI, but promotes brain tissue regeneration. **a** Representative images of TTC-stained brain sections from WT and HCAR1 KO mice 24 hours after HI. TTC turns red upon reacting with mitochondrial respiratory enzymes in viable tissue, whereas infarcted tissue remains white. **b** Brain infarct size (TTC-negative tissue as percentage of total quantified tissue volume) 24 hours after HI. **c** Percentage Infarct size per brain section 24 hours after HI. **d** Representative images of coronal brain sections from WT and HCAR1 KO mice 42 days after HI. **e** Brain tissue loss (% of total quantified tissue volume) 42 days after HI. **f** Percentage tissue loss per section 42 days after HI. Error bars indicate SD. Statistical significance was calculated using a two-tailed t-test (*p<0.05, **p<0.01, *** p<0.001, **** p<0.0001).

**Figure 2:**
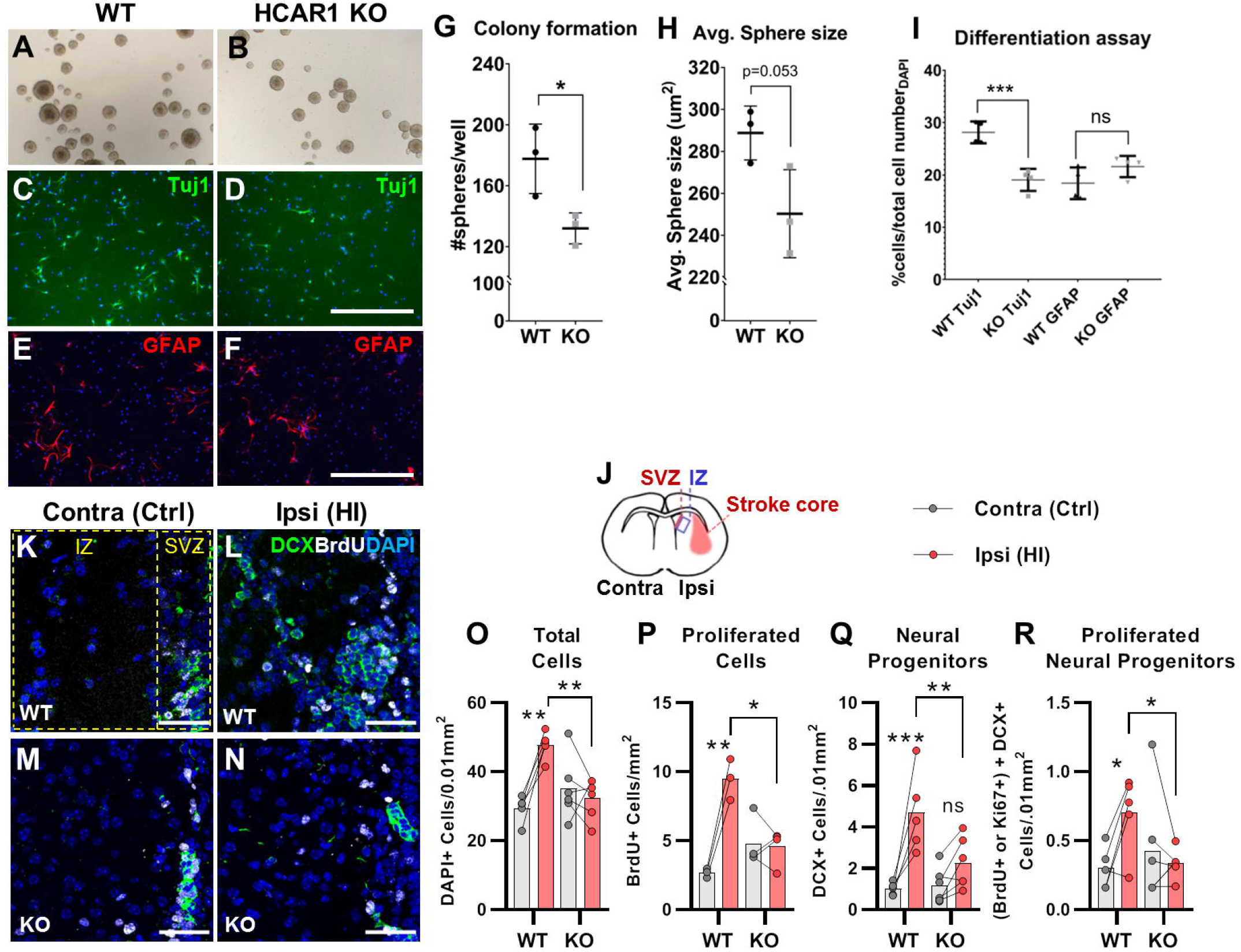
HCAR1 regulates neural stem and progenitor cell proliferation and differentiation. **a-i** Neurosphere formation from HCAR1 KO and WT cells. **a-b** Images of neurospheres from WT (a) and HCAR1 KO (b) mice. **c-f** Fluorescence images from WT (c, e) and HCAR1 KO (d, f) dissociated neurospheres after induced differentiation. Scale bare is 400 μm. **c-d** are stained with the neuronal marker Tuj1 and **e-f** are stained with the astrocyte marker GFAP. **g** Number of colonies formed per well. Each well was started with 10 000 cells. **h** Size of neurospheres (um^2^). **i** Percentage of cells positive for the neuronal marker Tuj1 or the astrocyte marker GFAP after induced differentiation of dissociated neurospheres. Data in **g-i** are shown as mean ±SD. n = 3 clones per genotype. **j-r** quantification of proliferating cells and neural progenitors after HI. **j** animation of a coronal mouse brain section illustrating the core of the infarct in the ipsilateral hemisphere (IPSI), the contralateral hemisphere (contra, used as control) as well as the proliferative subventricular sone (SVZ) and intermediate zone (IZ). **k-n** confocal images from coronal mouse brain sections labelled for DAPI (blue), doublecortin (DCX, marker of neuronal progenitor cells, green) and BrdU (injected proliferation marker, white). The images show the subventricular and intermediate zones in the contralateral (k, m) and ipsilateral (l, n) hemispheres in WT (k-l) and KO (m-n) mice. **o-q** Density of DAPI+ nuclei (i.e. all cells, o), BrdU+ cells (all proliferating cells, p) and DCX+ cells (neural progenitor cells, q) in the intermediate zones of the ipsi-(pink bars) and contralateral (white bars) hemispheres of WT and KO mice. r density of proliferating neural progenitor cells (i.e. cells that were both DCX+ and BrdU+ or Ki67+). *p<0.05; **p<0.01. Each point represents one sample/mouse. Ipsi-and contralateral samples from the same mouse are indicated with a line. WT n=3-5, KO n=4-6. Scale bars are 50 μm.

**Figure 3:**
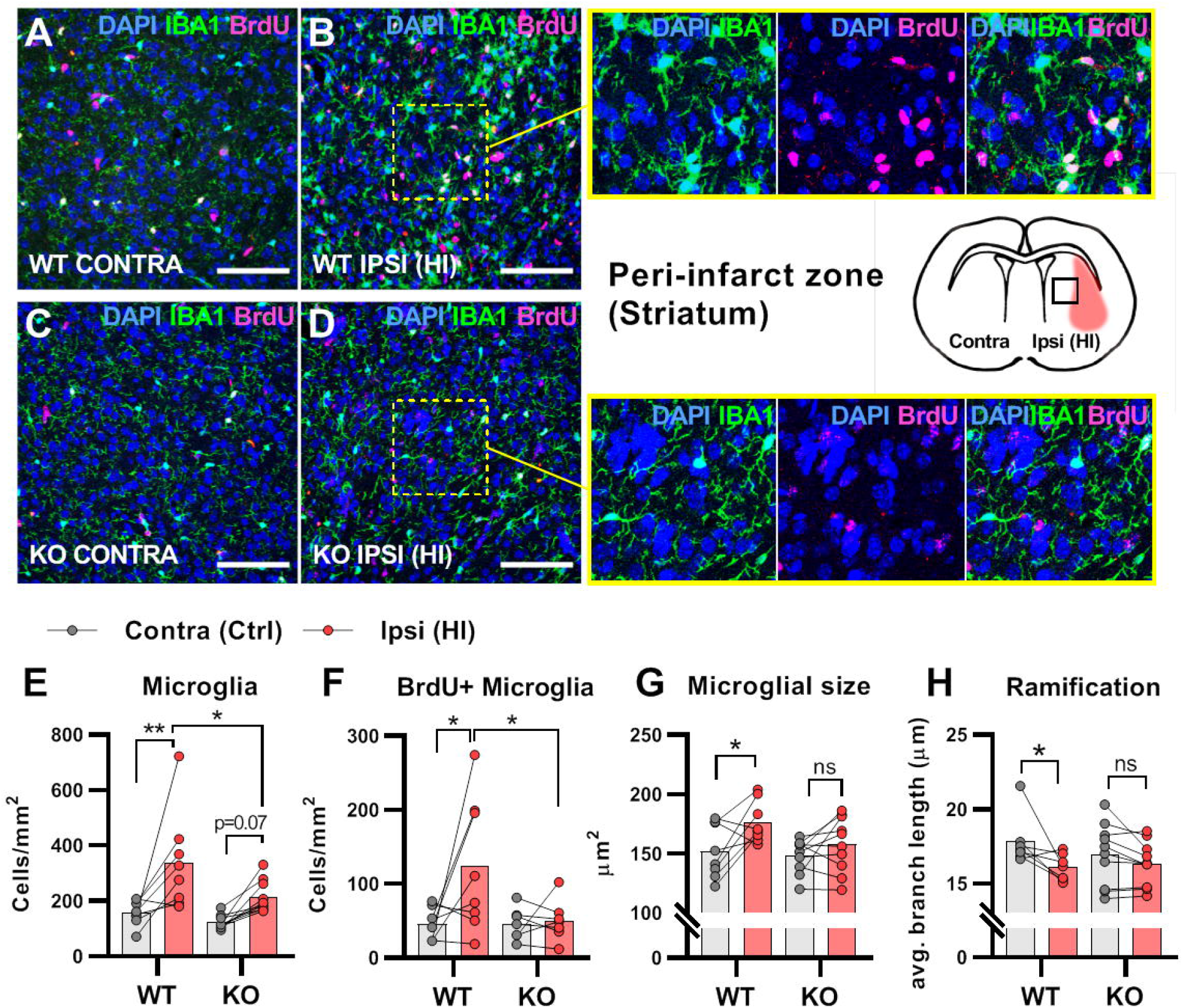
HCAR1 KO mice have deficient activation and proliferation of microglia after HI. **a-d** Confocal images from the peri-infarct zone (b, d, indicated as square in cartoon) and corresponding contralateral area (a, c) of coronal mouse brain sections from WT (a-b) and KO (c-d) labelled for BrdU (proliferating cells, magenta), Iba1 (microglia, green) and DAPI (blue nuclei). Scale bars are 100 μm. **e** Density of microglia (IBA1+ cells) in the peri-infarct zone (pink bars) and contralateral striatum (white bars) of WT and KO mice. **f** Density of proliferating microglia (i.e. cells that were both IBA1+ and BrdU+). **g-h** Assessment of microglia activation by morphology. When activated, microglia somata increase in size and get shorter and fewer branches. **g** average size of microglia somata. **h** Average maximum branch length. *p<0.05; **p<0.01. Each point represents one sample/mouse. Ipsi- and contralateral samples from the same mouse are indicated with a line. WT n=8, KO n=7-10.

### Data availability

The data that support the findings of this study as well as the algorithms used for analyses are available from the corresponding author upon request.

## Results

### HCAR1 is required for brain tissue regeneration after HI

To investigate the role of HCAR1 in stress-induced neuronal injury and subsequent neurogenesis, we induced HI in 9 days old HCAR1 KO and wild-type (WT) mice. We used a model for cerebral HI that includes permanent occlusion of the left common carotid artery followed by systemic hypoxia^14^. This leads to a detectable histological injury in the cortex, hippocampus, striatum, and thalamus of the left hemisphere, whereas the contralateral hemisphere is indistinguishable from a sham-treated brain, constituting a morphologically accurate internal control^14^. After HI, we examined acute brain tissue damage and long-term tissue loss. We assessed acute brain tissue damage by 2,3,5-Triphenyltetrazolium chloride (TTC) staining. HCAR1 KO and WT mice showed comprehensive damage in the affected ipsilateral hemisphere 24 hours after HI, with an average TTC-negative (i.e. damaged) volume in the ipsilateral relative to the contralateral side of 34 ± 22% in WT and 47 ± 14% in HCAR1 KO. There was no significant difference between HCAR1 KO and WT mice in total acute tissue damage (p=0.19, n=14 mice/genotype. Fig. 1A-B). When comparing the damage in individual sub-sections of the brain, we detected increased damage in HCAR1 KO mice in sections 4 mm from the frontal pole (WT 28 ± 27%; KO 46 ± 23%, p=0.046), but no difference between HCAR1 KO and WT in other sub-sections (Fig. 1C). Thus, overall there was little difference in acute tissue damage between HCAR1 KO and WT.

After the acute phase of HI, there is a phase of neurogenesis and tissue repair. To assess a potential role for HCAR1 in tissue repair, we measured tissue loss 42 days after HI. At this time point the repair process is completed and the long-term damage can be observed as a loss of brain tissue^14^. WT mice showed partial restoration of damaged brain structures with a tissue loss of 17 ± 9% (Fig. 1D-E). In comparison, HCAR1 KO mice showed significantly more tissue deficit with a permanent tissue loss of 31 ± 10%, i.e. 82% higher than in WT mice (Fig. 1D-E, p<0.0001 WT n=22, KO n=20). Measurements of tissue loss in individual sub-sections of the brain showed that all quantified sections (of which hippocampus, thalamus and striatum are included) had significantly lower regeneration in HCAR1 KO compared with WT (Fig. 1 F, 2 mm from pole WT 18 ± 10%, KO 27 ± 11%, p=0.007; 3 mm from pole WT 18 ± 9, KO 34 ± 12, p<0.0001, 4 mm from pole WT 18 ± 12, KO 35 ± 14, p=0.0001, 5 mm from pole WT 17 ± 11, KO 35 ± 12, p<0.0001, 6 mm from pole WT 14 ± 10, KO 31 ± 13, p<0.0001, 7 mm from pole WT 12 ± 7, KO 22 ± 10, p<0.007, n = at least 19 mice/genotype for 2-6 mm from pole and at least 10 mice/genotype for 7 mm from pole). In sum, mice lacking HCAR1 showed a significant deficit in regenerated tissue compared with WT mice, suggesting that HCAR1 is important for induced neurogenesis and tissue repair after HI.

### Impaired proliferation of neural progenitor cells in HCAR1 KO mice

Tissue repair after an ischemic injury is aided by an increase in proliferation and differentiation of neural stem- and progenitor cells (NSPCs)^15,16^. To test the effect of HCAR1 on NSPC proliferation and cell fate, we performed a neurosphere assay on spheres derived from forebrains of HCAR1 KO and WT mice. We found that neurospheres from HCAR1 KO mice developed fewer colonies compared with neurospheres from WT mice (Fig. 2A-B, G, no of colonies per well WT 178 ± 23; KO 132 ± 10, p = 0.034, df 4, n = 3 clones/genotype). The average size of HCAR1 KO spheres also tended to be smaller, although this was not statistically significant (Fig. 2A-B, H, sphere area WT 289 ± 13 μm^2^; KO 250 ± 21 μm^2^, p = 0.053, df 4, n = 3 clones/genotype). These results indicate that HCAR1 KO NSPCs have a lower self-renewal and proliferation rate. To examine the ability of the NSPCs to differentiate, the neurospheres were dissociated, and the NSPCs were cultured in differentiation medium for five days and immunolabelled for neurons and astrocytes. Both cell types were present after differentiation. However, HCAR1 KO cells had a lower percentage of Tuj1+ neurons compared with WT cells (Fig. 2 C-D, I, WT Tuj1 28 ± 2 %; KO Tuj1 19 ± 2 %, p < 0.001, df 12, n=4 clones/genotype), whereas the percentage of GFAP+ astrocytes was not significantly different between the two genotypes (Fig. 2 E-F, I, WT GFAP 18 ± 3 %; KO GFAP 22 ± 3 %, p = 0.08, df 12, n= 4 clones/genotype). This suggests that HCAR1 directs stem cells towards a neuronal fate.

As the neurosphere data suggested impaired proliferation of NSPCs, we then examined *in vivo* cell proliferation after HI in HCAR1 KO and WT mice. Mice were injected with the proliferation marker Bromodeoxyuridine (BrdU) on days 4-7 after HI and were sacrificed on day 7. The number of BrdU+ cells was assessed by immunohistochemistry on brain sections. We focused on the subventricular zone and the adjacent intermediate zone as these are proliferative areas that contain a large portion of neuronal progenitor cells^4^. In the intermediate zone, the total number of cells was increased by 63% in the affected ipsilateral hemisphere compared with the contralateral hemisphere in WT mice (Fig. 2K-O, DAPI+ cells/0.01 mm^2^: contra 29.4 ± 3.8, ipsi 47.8 ± 4.0, p=0.001, df 9, n=5). WT mice also had a significant increase in the number of newly proliferated BrdU+ cells on the ipsilateral side (Fig. 2K-N, P, BrdU+ cells/0.01 mm^2^: contra 2.7 ± 0.3, ipsi 9.5 ± 1.5, p=0.001, df 9, n=3). In KO mice, however, there was no change in the total number of cells or the number of newly proliferated cells in the ipsilateral compared with contralateral side Fig. 2K-P, KO DAPI+ cells/0.01 mm^2^: contra 35.1 ± 9.0, ipsi 32.45 ± 5.8, p=0.66, df 9, n=6. BrdU+ cells/0.01 mm^2^: contra 4.75 ± 1.8, ipsi 4.6 ± 1.3, p=0.97, df 9, n=4). Thus, in line with the neurosphere data, it appears that HCAR1 KO mice are unable to induce cell proliferation after HI. We then looked at neural progenitor cell proliferation. The number of neural progenitor cells positive for the marker doublecortin (DCX) were strongly increased in the ipsilateral side after HI in WT, but did not have a statistically significant increase in HCAR1 KO mice (Fig. 2K-N, Q. DCX+ cells/0.01mm^2^: WT contra 1.0 ± 0.3, ipsi 4.7 ± 2.0, p=0.001, df 9, n=5. KO contra 1.2 ± 0.9, ipsi 2.3 ± 1.2, p=0.21, df 9, n=6). Further, the number of newly proliferated DCX+ cells increased on the ipsilateral side in WT but not in KO (Fig. 2K-N, R. DCX+ and Ki67+ (n=2) or BrdU+ (n=3 WT and 4 KO) cells/0.01 mm^2^: WT contra 0.30 ± 0.15, ipsi 0.70 ± 0.28, p=0.03, df 9. KO contra 0.43 ± 0.40, ipsi 0.34 ± 0.10, p=0.74, df 9, n=5 WT and 6 KO). A similar trend with fewer progenitor cells after HI in HCAR1 KO compared with WT was seen in the subventricular zone, although here there was large variation between samples, possibly due to the small size of this region (Supplementary Fig. 1). In conclusion, these data show that HCAR1 KO mice fail to increase proliferation of neural progenitor cells after HI, suggesting that HCAR1 is required to stimulate neural cell proliferation to induce tissue repair after ischemic damage.

### Impaired microglial proliferation and activation in HCAR1 KO mice after HI

After cerebral HI, microglia are recruited to the injured site at which they remove debris from dead cells to facilitate the repair process^17^. This requires increased proliferation, activation and migration of the microglia^17,18^. We used immunohistochemistry to assess the proliferation and activation of microglia in the area surrounding the infarct (the peri-infarct zone). As expected, WT mice had a strong increase in proliferating microglia in the ipsilateral hemisphere when compared with the contralateral side. However, no increase was detected in HCAR1 KO mice (Fig. 3A-D, F. IBA1+ BrdU+ cells/mm^2^: WT contra 45.4 ± 21.7, ipsi 123.2 ± 90.0, p=0.01, df 26, n=8. KO contra 45.5 ± 21.5, ipsi 50.4 ± 27.8, p=0.98, df 26, n=7.). Further, the total number of microglia was increased in the ipsilateral side in WT, but not significantly in HCAR1 KO mice (Fig. 3A-D, E. IBA1+ cells/mm^2^: WT contra 156.0 ± 43.3, ipsi 337.5 ± 178.2, p=0.002, df 16, n=8. KO contra 121.8 ± 57.0, ipsi 214.0 ± 57.0, p=0.074, df 16, n=10.). We then assessed the activation of microglia. Activated microglia have larger cell soma and are less ramified (i.e. they have shorter and fewer branches)^19^. In WT mice, microglia in the ipsilateral hemisphere had larger somata and were less ramified than in the contralateral side, indicating an activated phenotype (Fig. 3A-B, G-H. Cross-sectional area, μm^2^: WT contra 152.0 ± 22.8, ipsi 176.5 ±17.5, p=0.02, df 16, n=8. Ramification (avg. max branch length, μm): WT contra 17.8.1 ± 1.7, ipsi 16.1 ±0.9, p=0.01, df 15. n=7. Surprisingly, no significant changes in the cell soma size and ramification of microglia were observed in HCAR1 KO mice (Fig. 3C-D, G-H, cross-sectional area, μm^2^: KO contra 148.1 ± 14.0, ipsi 158.2 ±23.3, p=0.37, df 16, n=10. Ramification (Avg. max branch length, μm): KO contra 17.0 ± 2.1, ipsi 16.3 ±1.5, p=0.36, df 15, n=10), indicating that microglia activation is not induced by HI. In sum, HCAR1 KO mice were unable to induce microglia proliferation and activation in response to HI, indicating a role for HCAR1 in these processes.

### Weak transcriptional response to HI in the subventricular zone of HCAR1 KO mice

To investigate the mechanisms underlying HCAR1 involvement in brain tissue regeneration, we performed a genome-wide transcriptome analysis by RNA sequencing of the subventricular region from the affected ipsilateral and contralateral (control) hemispheres of mice after cerebral HI. Principal component analysis (PCA, Fig. 4A) showed that tissue samples from the ipsilateral hemisphere of WT mice clustered away from the contralateral samples (i.e. they showed a different gene expression profile), indicating a strong transcriptional response to HI. Samples from the contralateral hemisphere of HCAR1 KO mice had a comparable gene expression pattern to WT contralateral samples. Notably, ipsilateral HCAR1 KO samples also clustered close together with WT and KO contralateral samples. Thus, it appears that the transcriptional response to HI in the subventricular zone of HCAR1 KO is severely impaired. The number of differentially expressed genes (DEGs) between the different experimental groups further reflected an inadequate response in HCAR1 KO (Supplementary table 1): while the WT ipsilateral hemisphere had 7332 DEGs when compared with WT contralateral hemisphere, indicating a distinct response to HI, only 752 DEGs were detected between HCAR1 KO ipsilateral and contralateral hemispheres. Further, when comparing WT contralateral with HCAR1 KO contralateral hemisphere, only 11 DEGs were identified, whereas WT ipsilateral versus KO ipsilateral identified 6640 DEGs. Therefore, in the undamaged contralateral side, WT and HCAR1 KO showed a similar gene expression profile, while HI induced a large gene expression response in WT that was strongly reduced in HCAR1 KO. To investigate whether the deficient transcriptional response to HI was specific to the subventricular zone, we performed a similar RNA sequencing analysis of the ipsilateral and contralateral hippocampi from the same mice. Surprisingly, in the hippocampal samples, PCA analysis showed a close clustering of HCAR1 KO and WT samples after HI (Supplementary Fig. 6), indicating a similar transcriptional response to HI. In line with this, we only identified 37 DEGs when comparing HCAR1 KO with WT hippocampi after HI (Supplementary table 1). This indicates that HCAR1 acts as a key transcriptional regulator of ischemia response in the subventricular zone but not in the hippocampus.

**Figure 4:**
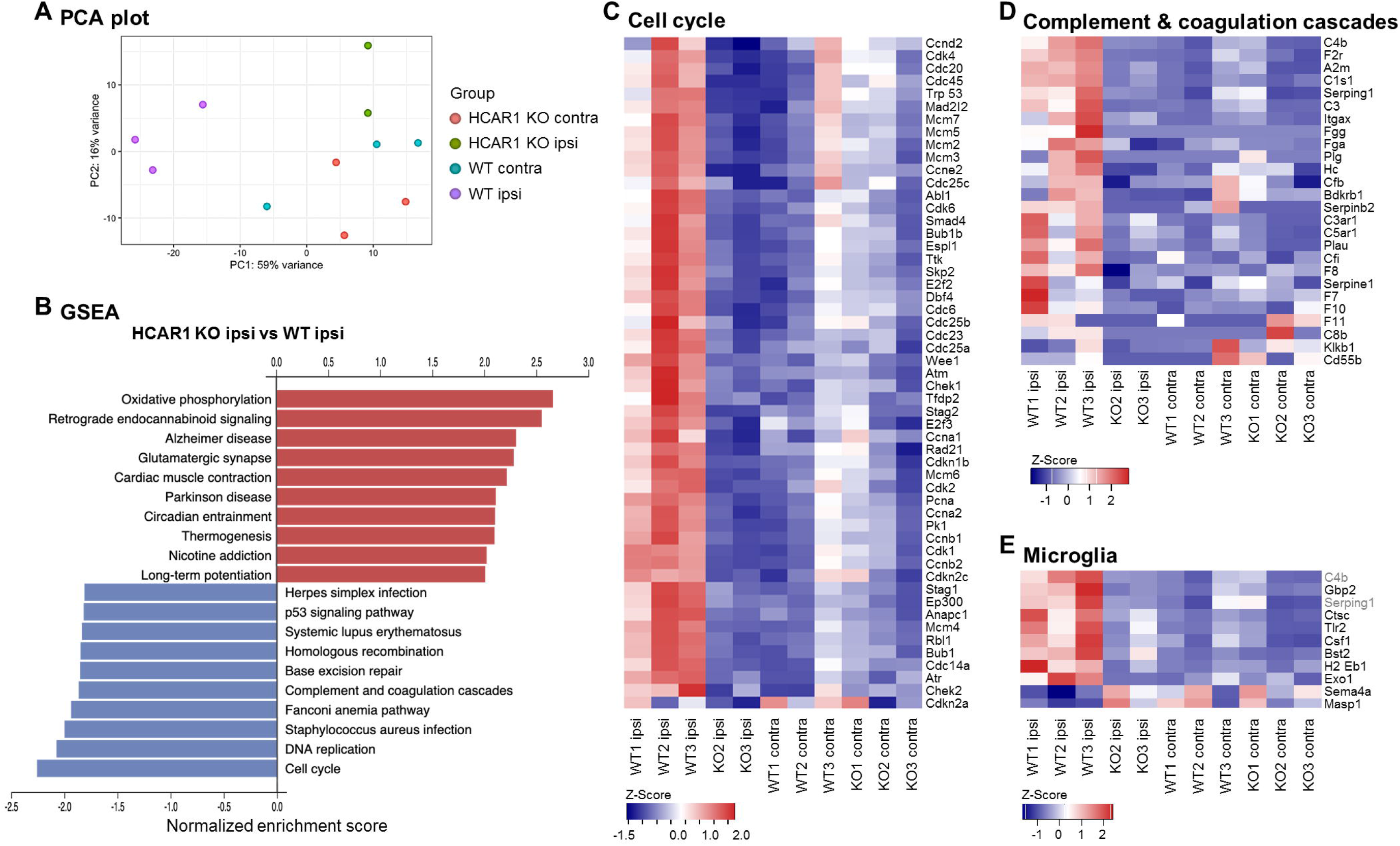
HCAR1 regulates transcriptional response to ischemia including cell cycle and complement pathway. **a** PCA plot of transcriptome data from subventricular zone tissue samples from the ipsilateral (HI-damaged) and contralateral (control) hemisphere in HCAR1 KO and WT mice. Each point represents one sample/mouse. Each colour represents a group. **b** Gene set enrichment analysis (GSEA) of HCAR1 KO ipsi versus WT ipsi showing the ten most up- or downregulated pathways (FDR<0.05). **c** Heatmap showing relative expression of a subset of differentially expressed genes (DEGs) enriched in the cell cycle. **d** DEGs related to complement and coagulation cascades (immune system response involved in activation of microglia. **e** DEGs that are markers of microglia activation (based on DePaula-Silva et al., 2019^42^). Genes shown in c-e were differentially expressed in HCAR1 KO ipsi versus WT ipsi, but expression is shown in all samples from all four experimental groups (se bottom of figures).

We then performed gene set enrichment analysis of the subventricular zone samples to identify differentially regulated pathways between the two genotypes in this area. Several pathways were differentially expressed in HCAR1 KO ipsi versus WT ipsi (Fig. 4B. For extensive maps of differentially regulated pathways between all experimental groups, see Supplementary figures 2-5). Of particular interest to our previous findings, we found the cell cycle pathway strongly down-regulated in HCAR1 KO. Furthermore, the P53 pathway, a well-known regulator of the cell cycle, was also down-regulated. The relative expression of differentially regulated cell cycle genes across the four different experimental groups is shown in Fig. 4C. The downregulation of cell cycle genes in HCAR1 KO compared with WT may explain the differences in neural differentiation and the deficient cell proliferation in HCAR1 KO after HI (Fig. 2).

The complement and coagulation cascade pathways were down-regulated in the ipsilateral hemisphere of HCAR1 KO (Fig. 4B, D) compared with WT mice. This is of particular interest in light of the diminished microglia response in HCAR1 KO as the complement system is involved in microglia activation^20,21^. A high number of markers for activated microglia were also down-regulated in HCAR1 KO vs WT after HI (Fig. 4E), also in line with the impaired microglia activation observed by immunostaining (Fig. 3). In sum, the subventricular zones of HCAR1 KO mice display a strongly impaired transcriptional response to HI. This can explain the impaired cell proliferation and microglia activation after HI, suggesting that HCAR1 is a key transcriptional regulator of tissue repair after ischemia.

## Discussion

We report that HCAR1 KO mice have a substantial deficit in the restoration of brain tissue after HI, indicating that lactate receptor HCAR1 plays a crucial role in the processes that lead to tissue repair. Since no exogenous lactate was administered in our experiments, the observed effect must be due to endogenous lactate or possible baseline receptor activity. Indeed, the lactate level rises after a hypoxic-ischemic episode^22,23^. It is likely that administration of lactate could further leverage the effect of HCAR1: two recent studies showed that mouse pups injected with lactate before, or in the hours or days following HI had improved recovery^7,8^. The authors suggested that the protective effect of lactate was mainly due to lactate being used as a metabolite to make ATP^7,8^. However, our data suggest that recovery after lactate injection is partly mediated by HCAR1, which promotes a stronger transcriptional response to HI and thereby facilitates neurogenesis and tissue regeneration after injury. On the other hand, these studies also showed an effect of lactate injections on acute infarct volume. Therefore, a putative explanation is that lactate injected before or immediately after HI reduces lesion size by mainly working as a metabolite, whereas lactate injected at a later time point in large works via HCAR1. Lactate injections can also be protective after ischemic stroke in adult mice^24–27^. Here, it also seems to be a combination of metabolic and HCAR1-dependent effects since replacing lactate with either the HCAR1 receptor agonist 3,5-dihydroxybenzoic acid (3, 5 DHBA) or the metabolic substrate pyruvate offered partial protection^24^.

An ischemic event will induce a significant transcriptional response^28^. By RNA sequencing of tissue samples from the subventricular zone, we found that HCAR1 KO mice displayed a weak transcriptional response to HI, with a 90% reduction in DEGs compared with WT mice. Hence, HCAR1 appears to be essential for the induction of the transcriptional response to HI. This HCAR1 dependence is specific for the subventricular zone, since hippocampal samples showed very moderate differences between WT and HCAR1 KO mice in the transcriptional profile after HI.

An essential part of the repair process after a neonatal brain injury is the generation of new cells. This occurs by increased proliferation and differentiation of stem cells and involves upregulation of genes involved in the cell cycle pathway^29,30^. By use of neurosphere assays, we showed that neurospheres lacking HCAR1 had reduced proliferation ability. Moreover, while WT mice more than doubled the number of proliferating cells after HI, HCAR1 KO mice were unable to increase cell proliferation (Fig. 2P). In line with this, transcriptome analysis revealed that the cell cycle genes were strongly upregulated after HI in WT, but not in HCAR1 KO mice. Hence, it seems that HCAR1 can act as a transcriptional regulator of cell cycle genes, thereby controlling cell proliferation. A role of HCAR1 in cell proliferation was previously shown in cancer and osteoblast cell lines^12,13^. A very recent paper also showed HCAR1 dependent neurogenesis after high intensity physical exercise^43^. Similar to our study, this paper found neurogenesis to be HCAR1 dependent in the subventricular zone, but not in the hippocampus. Under normal conditions, however, HCAR1 may be less important as we did not detect any differences between HCAR1 KO and WT mice in the total number of cells, or proliferating cells, in the contralateral hemispheres. This was evident from the transcriptome analysis of contralateral tissue showing only moderate differences in DEGs between KO and WT.

Microglia carry out several vital functions in response to brain injury. These include clearance of damaged tissue, resistance to infections and restoration of tissue homeostasis^5^. We detected an increase in the proliferation and activation of microglia in the peri-infarct zone after HI in WT, but not in HCAR1 KO mice. The lack of microglia proliferation could be due to the reduced induction of cell cycle response, as discussed above. The absence of microglia activation was confirmed on a transcriptional level as genes associated with activated microglia were upregulated in WT but not in KO. This effect may be explained by the reduced complement system response in HCAR1 KO. Altogether, the differences in microglial data suggest that the damaging effects of HCAR1 KO in HI are due to interplay of both the immune response and proliferation and regeneration of the damaged brain cells.

In addition to the effects on cell cycle and microglia activation discussed above, the transcriptome analysis revealed a large number of differentially expressed genes and pathways, including genes involved in DNA repair and glutamate signalling. These processes will likely also influence the ability of the brain tissue to repair after injury.

Overall, our data show that HCAR1 is a key transcriptional regulator of brain tissue response to an ischemic insult. We therefore propose a model in which activation of HCAR1 by elevated lactate during and after HI stimulates a transcriptional response involving pathways responsible for tissue repair (Fig. 5). HCAR1 could be a target of future treatment for neonatal HI and possibly other forms of brain injury.

**Figure 5:**
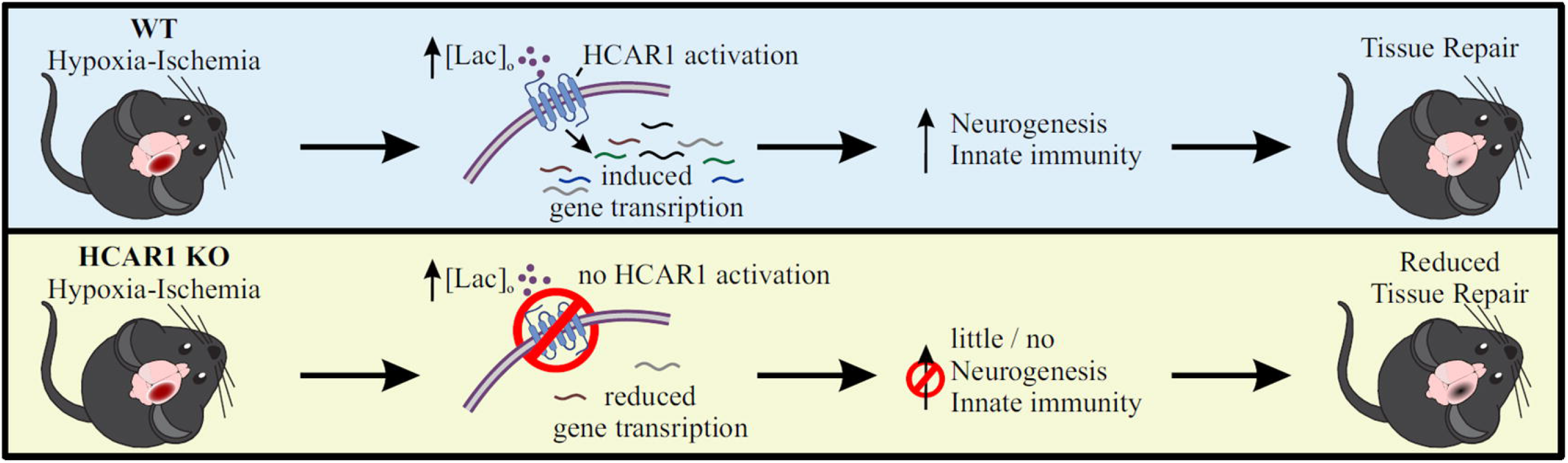
Proposed model for the role of HCAR1 in neonatal HI. During and after HI, the extracellular concentration of lactate ([lac]_o_) is elevated. **Top panel:** In WT mice, the elevated lactate causes HCAR1 activation, which induces transcription of genes involved in tissue response to ischemia. This includes genes responsible for neurogenesis and innate immunity, thereby promoting tissue repair. **Bottom panel:** In HCAR1 KO mice, the transcriptional response to ischemia is severely reduced. Without the HCAR1-induced gene transcription, there is little neurogenesis and innate immune response, which in turn gives an impaired tissue repair.

## Acknowledgements

We thank Prof. Stefan Offermanns and collaborators at the Max-Planck-Institute for Heart and Lung Research, Bad Nauheim, Germany, for providing breeder HCAR1 knockout mice, Dr. Gunn Anette Hildrestrand for assistance with hypoxic-ischemic experiments and Dr. Adam Filipczyk for comments on the manuscript. Images were obtained with support from Drs. Anna Lång and Stig-Ove Bøe at the South-Eastern Health Authority Core Facility of Advanced Light Microscopy (Gaustad, Norway).

## Supplementary material

All supplementary material is uploaded in a separate file.

## Author contributions

L.H.K., E.R.G., V.P., M.P., W.W., A.A-J., R.S., N.M., X.L. and J.E.R. performed and analysed experiments. L.H.K., E.R.G., V.P., L.H.B., M.B. and J.E.R. planned experiments and wrote the manuscript.

