## Supplementary material for "Lactate receptor HCAR1 regulates neurogenesis and microglia activation after neonatal hypoxia-ischemia"

### Supplementary information

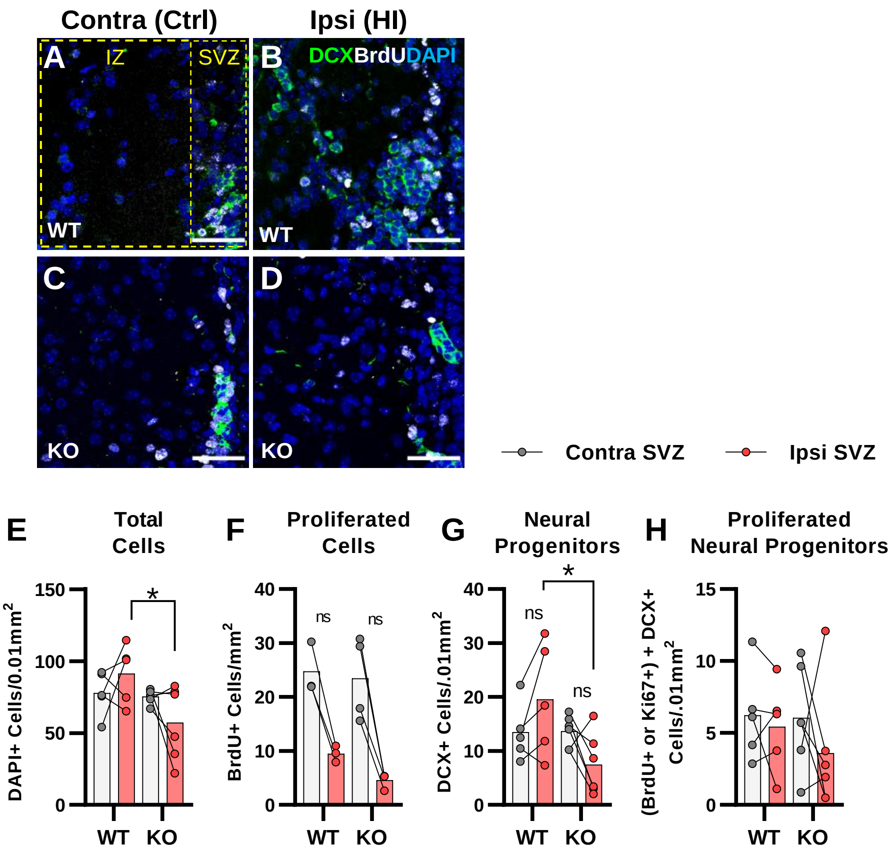


**Supplementary figure 1. Quantification of proliferating cells and neural progenitors in the subventricular zone after HI. a-d** confocal images from coronal mouse brain sections labeled for DAPI (blue), doublecortin (DCX, marker of neuronal progenitor cells, green) and BrdU (injected proliferation marker, white). The images show the subventricular and intermediate zones in the contralateral (a, c) and ipsilateral (b, d) hemispheres in WT (a-b) and KO (c-d) mice. **e-g** Density of DAPI+ nuclei (i.e. all cells, e), BrdU+ cells (all proliferated cells, f) and DCX+ cells (neural progenitor cells, g) in the subventricular zones of the ipsi- (pink bars) and contralateral (white bars) hemispheres of WT and KO mice. **h** density of proliferated neural progenitor cells (i.e. cells that were both DCX+ and BrdU+ or Ki67+). *$p<0.05$. The paired points with line represent n per genotype (WT n=3-5, KO n=4-6). Scale bars are 50 µm.

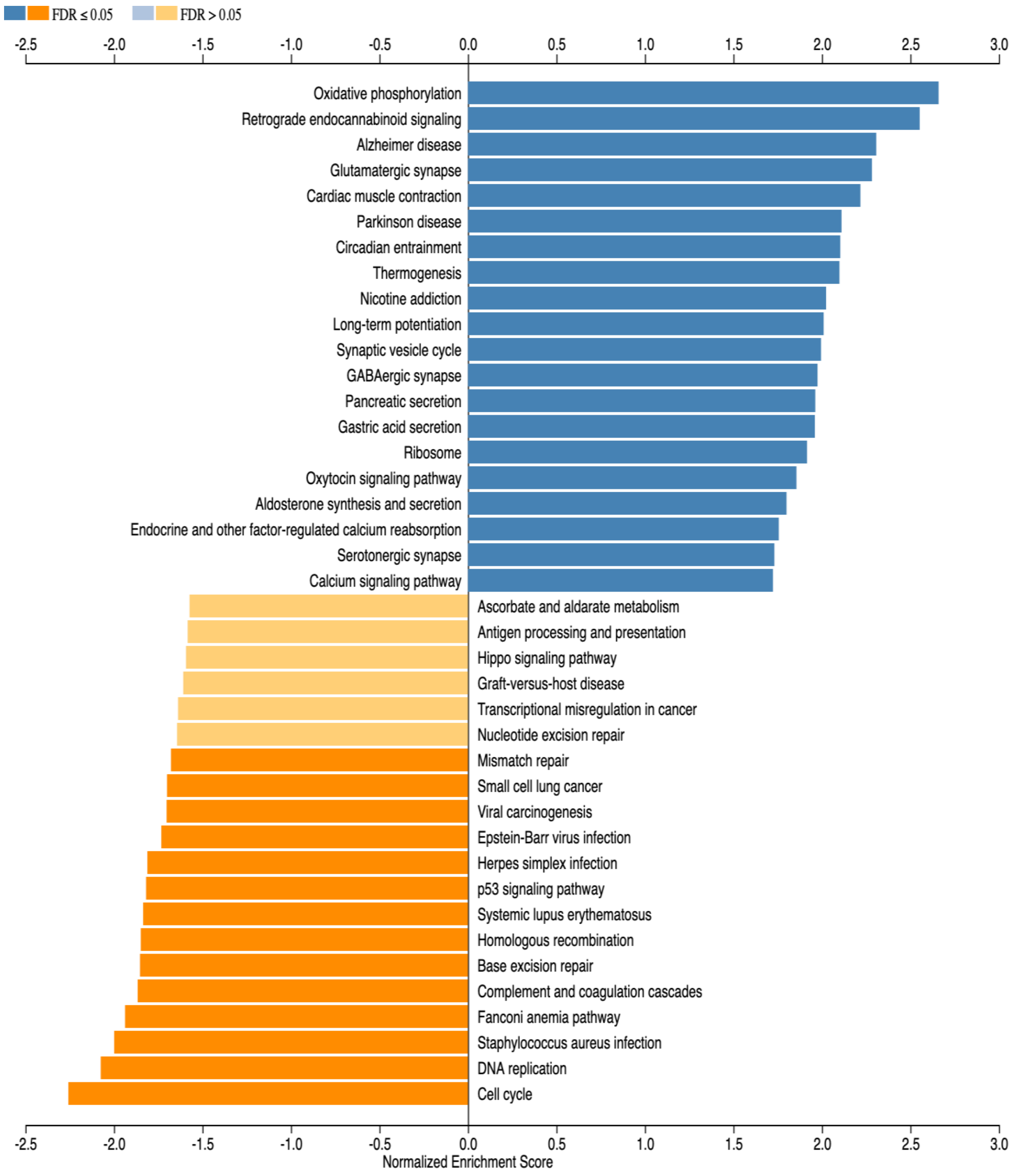


**Supplementary figure 2. Gene set enrichment analysis (GSEA) of subventricular zone from HCAR1 KO ipsi versus WT ipsi.** The 20 most up- or downregulated pathways are included.

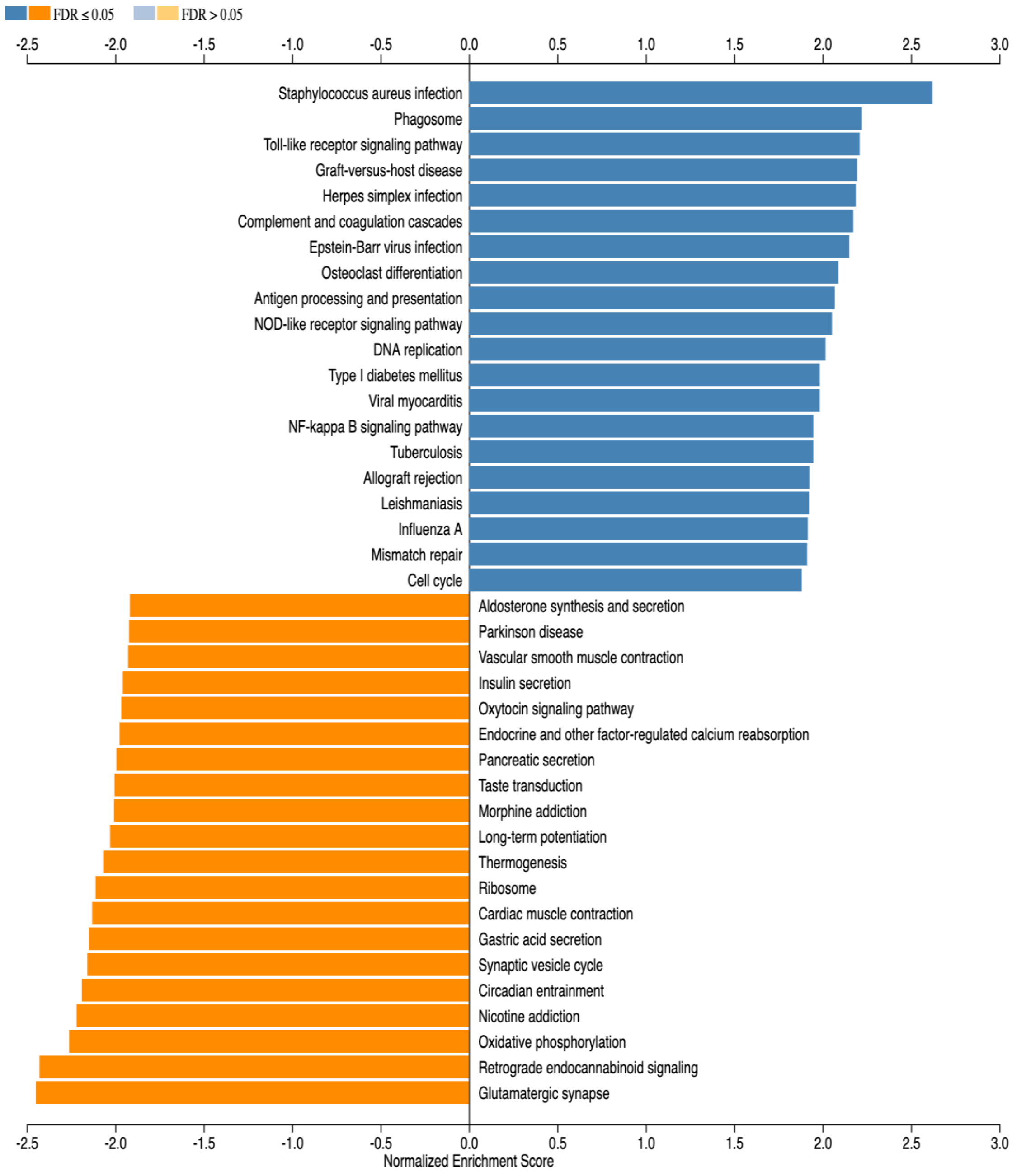


**Supplementary figure 3. Gene set enrichment analysis (GSEA) of subventricular zone from WT ipsi (HI) versus WT contra (control).** The 20 most up- or downregulated pathways are included.

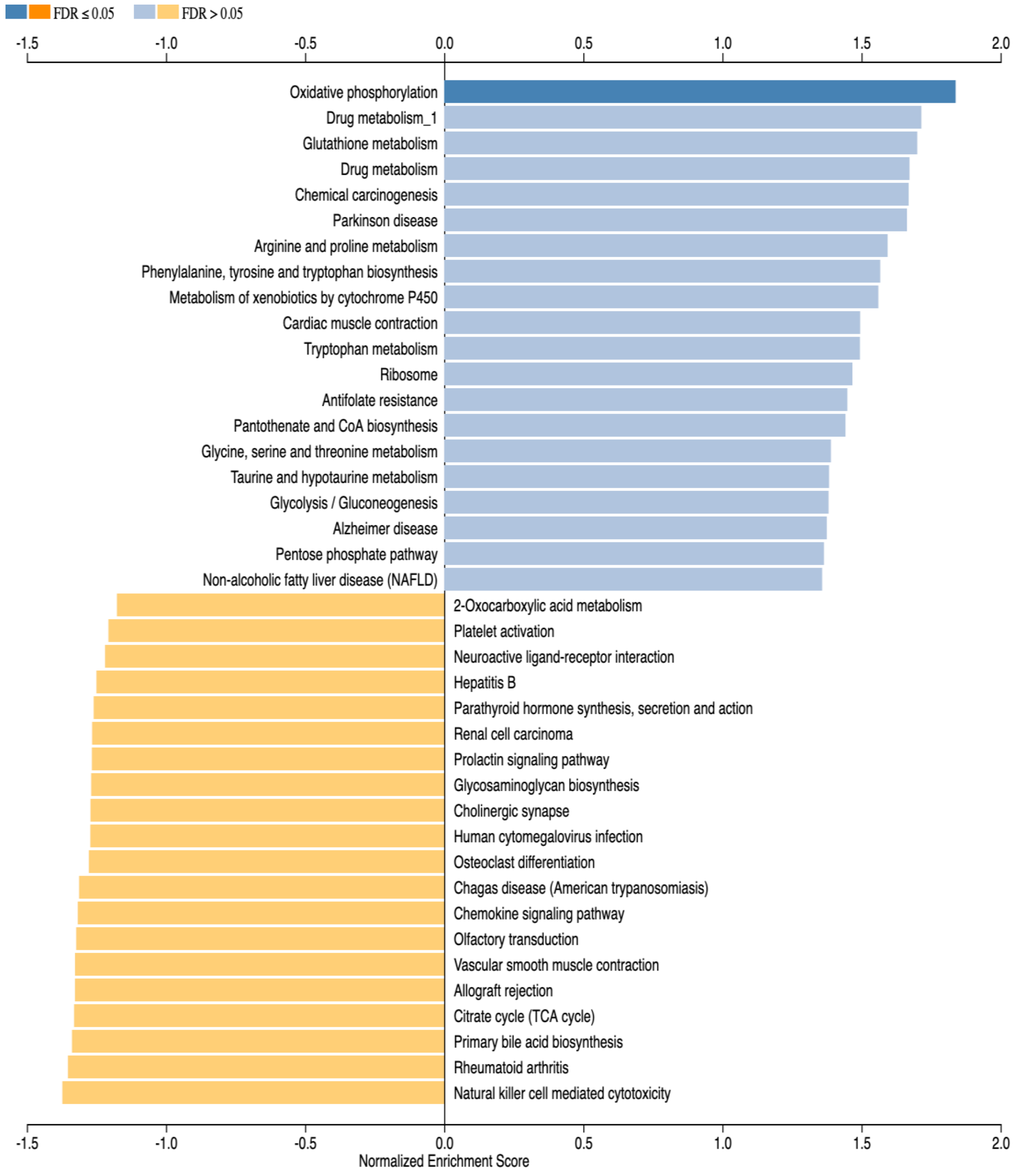


**Supplementary figure 4. Gene set enrichment analysis (GSEA) of subventricular zone from HCAR1 KO contra versus WT contra.** The 20 most up- or downregulated pathways are included. Note that only one pathway has FDR≤0.05.

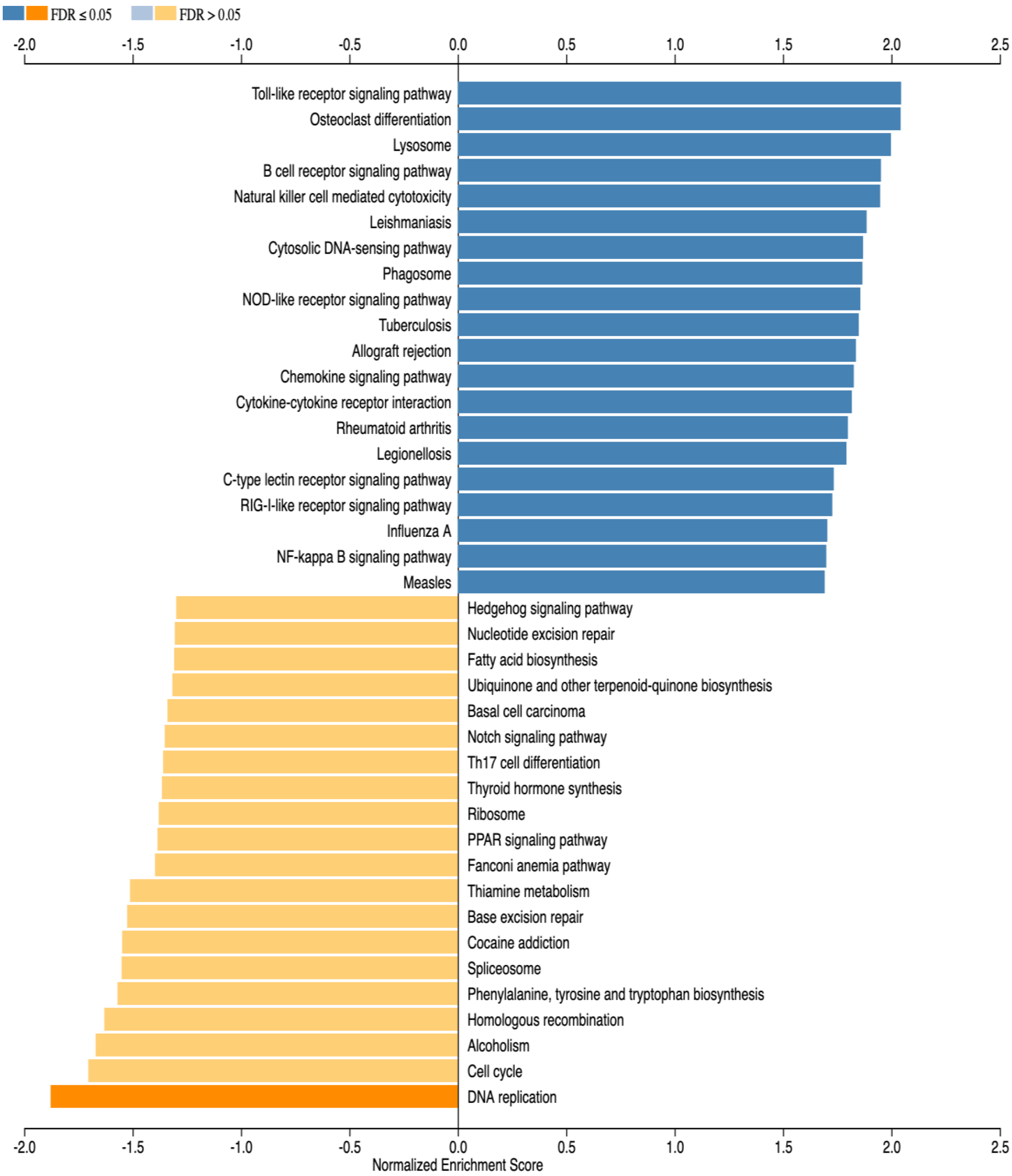


**Supplementary figure 5. Gene set enrichment analysis (GSEA) of subventricular zone from HCAR1 KO ipsi (HI) versus HCAR1 KO contra (control).** The 20 most up- or downregulated pathways are included.

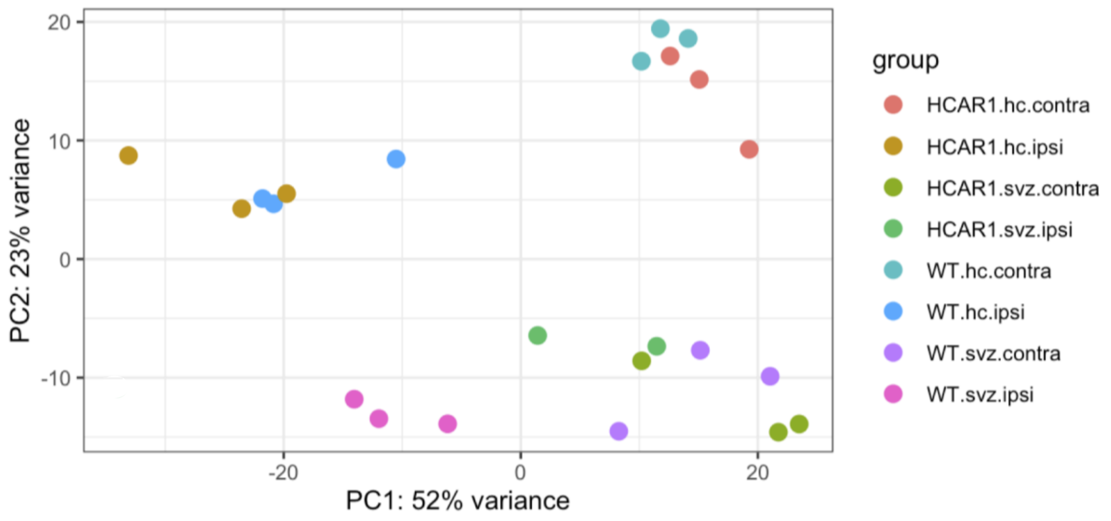


**Supplementary figure 6. HCAR1 KO mice show a transcriptional response to HI in the hippocampus, but not in the subventricular zone.** PCA plot of transcriptome data from hippocampal and subventricular zone tissue samples from the ipsilateral (HI-damaged) and contralateral (control) hemisphere in HCAR1 KO and WT mice. Each point represents one sample/mouse. Each color represents a group. Note that the data from subventricular zone samples are the same as shown in Figure 4 in the main text.

**Supplementary table 1. Number of differentially expressed genes (DEGs) between the different experimental groups**

| Comparisons | Total UP DEGs | Total DOWN DEGs | Total DEGs |
|---|---|---|---|
| HCAR1.svz.ipsi vs WT.svz.ipsi | 3258 | 3182 | 6440 |
| WT.svz.ipsi vs WT.svz.contra | 3594 | 3738 | 7332 |
| HCAR1.svz.contra vs WT.svz.contra | 8 | 3 | 11 |
| HCAR1.svz.ipsi vs HCAR1.svz.contra | 251 | 501 | 752 |
| HCAR1.hc.ipsi vs WT.hc.ipsi | 33 | 4 | 37 |
| WT.hc.ipsi vs WT.hc.contra | 2894 | 2298 | 5192 |
| HCAR1.hc.contra vs WT.hc.contra | 10 | 3 | 13 |
| HCAR1.hc.ipsi vs HCAR1.hc.contra | 4167 | 3625 | 7792 |

*Thresholds -> logFC ≥ 0.5 (UP DEGs) and ≤ -0.5 (DOWN DEGs); all of them significant at adjusted p-value < 0.05*
